# Cause and chondroprotective effects of prostaglandin E2 secretion during mesenchymal stromal cell chondrogenesis

**DOI:** 10.1101/2024.01.20.576230

**Authors:** S. Schmidt, F.A.M. Klampfleuthner, T. Renkawitz, S. Diederichs

## Abstract

Mesenchymal stromal cells (MSCs) that are promising for cartilage tissue engineering secrete high amounts of prostaglandin E2 (PGE2), an immunoactive mediator involved in endochondral bone development. This study aimed to identify drivers of PGE2 and its role in the inadvertent MSC misdifferentiation into hypertrophic chondrocytes. PGE2 release which rose in the first three weeks of MSC chondrogenesis was jointly stimulated by endogenous BMP, WNT, and hedgehog activity that supported the exogenous stimulation by TGF-β1 and insulin, and overcame the PGE2 inhibition by dexamethasone. Experiments with PGE2 treatment or the inhibitor celecoxib or specific receptor antagonists demonstrated that although driven by prohypertrophic signals, PGE2 exerted broad autocrine antihypertrophic effects. This chondroprotective effect makes PGE2 not only a promising option for future combinatorial approaches to direct MSC tissue engineering approaches into chondral instead of endochondral development, but could potentially have implications for the use of COX-2-selective inhibitors in osteoarthritis pain management.

## Introduction

Mesenchymal stromal cells (MSCs) are promising for the regeneration of musculoskeletal tissues, including cartilage, because they are a source of adult stem cells with multilineage differentiation capacity that is easily accessible from regenerative tissues such as bone marrow. Cartilage defects, that have been found in the knees of about 70% of patients without a prior history of osteoarthritis by high-resolution magnetic resonance imaging [1], are a frequent clinical problem. The restricted intrinsic repair capacity of articular cartilage makes surgical intervention often necessary to relieve the patient’s pain and disability and to minimize the risk of joint-wide cartilage degeneration and osteoarthritis. MSCs are an attractive alternative to autologous chondrocyte implantation (ACI) that is the current gold standard for treating focal chondral defects above 2cm^2^ in size [2]. However, ACI involves creating new cartilage lesions for cell harvest, again risking insufficient healing and osteoarthritis development. Unfortunately, when subjected in vitro to chondrogenic conditions including the chondro-inducing transforming growth factor-beta (TGF-β), MSCs inevitably undergo endochondral rather than chondral differentiation and develop a hypertrophic chondrocyte phenotype [3]. By contrast, articular chondrocytes stably remain in the chondral lineage under the same conditions. This misdifferentiation of MSCs is highly undesirable for cartilage repair because it predestines the resulting tissue engineering constructs to mineralization and endochondral bone formation in vivo [4]. MSC misdifferentiation thus recapitulates the normal development of growth plate chondrocytes and importantly, also the pathological degeneration of osteoarthritic articular chondrocytes. They all have in common the characteristic expression of the prohypertrophic transcription factor myocyte-specific enhancer factor 2C (*MEF2C*) along with calcifying proteins like type X collagen (*COL10A1*), alkaline phosphatase (*ALPL*), integrin-binding sialoprotein (*IBSP*), and secreted phosphoprotein type 1 (*SPP1*). Decades of comparative research with articular chondrocytes have identified that MSC misdifferentiation is driven by misregulated parathyroid hormone-related protein (PTHrP)/indian hedgehog (IHH) signaling, endogenous wingless-int (WNT) signaling [5–7], and putatively also TGF-β-receptor (ALK4/5)-mediated SMAD1/5/9 signaling (manuscript in review). Importantly, all attempts at reprogramming MSC chondrogenesis into stable chondral development in vitro by inhibiting the known prohypertrophic signaling pathways remained insufficient [6, 8, 9], suggesting that the drivers of endochondral misdifferentiation are still incompletely understood.

Interestingly, we recently found a strongly elevated expression of the cyclooxygenase*-*2 (COX-2)-encoding gene prostaglandin-endoperoxide synthase 2 (*PTGS2*) along with its enzymatic product prostaglandin E2 (PGE2) throughout MSC chondrogenesis compared to articular chondrocyte cultures[10], which raised our interest in a potential contribution of PGE2 to endochondral misdifferentiation. PGE2 signals through one of the four G-protein-coupled prostaglandin E2 receptors (EP) 1, 2, 3, or 4 and is best described for its broad regulation of various immune cells including dendritic cells and T cells (reviewed by [11]). Released by chondrocytes and other joint cells, PGE2 is also involved in osteoarthritis pathology as a regulator of inflammation, angiogenesis, osteoclast activation, and osteophyte formation [12–15], thus PGE2 inhibition may be warranted when implanting MSC-based engineered cartilage replacement tissue with high PGE2 release. However, the importance of PGE2 for MSC chondrogenesis and whether its inhibition would be beneficial or detrimental for the tissue engineered neocartilage, remains unaddressed.

The drivers of COX-2/PGE2 during MSC chondrogenesis beyond mechanical load[10] are currently unknown. Interestingly, in cells of various types and species including murine chondrocytes, TGF-β/BMP and WNT5A have been reported to stimulate COX-2/PGE2 production [16–19], suggesting that PGE2 production is a consequence of prohypertrophic signaling in differentiating MSCs. Beyond that, PGE2 has been functionally implicated in endochondral development by in vitro studies with the murine pro-chondrogenic cell line ATDC5 and rat periosteum cultures, in which COX-2 inhibition was reported to reduce *Col10a1* and *Alpl* gene expression [20], while in turn PGE2 stimulated *Col10a1* expression [21]. Moreover, an impaired endochondral bone formation and fracture healing was apparent in COX-2 loss-of-function models in mice, rats, and rabbits that could be rescued by PGE2 or an EP4 agonist [22–27]. While altogether, this may indicate a prohypertrophic activity of PGE2, a large body of opposing studies reported that PGE2 or EP4 activation decreased *Col10a1* and *Alpl* expression, ALP activity, and mineralization in chick or murine limb bud mesenchymal cells and growth plate chondrocytes [28–32], attributing an antihypertrophic activity to PGE2. Thus, while overall PGE2 has been implicated to have an active functional role in endochondral development by numerous studies in a wide range of biological systems, the exact nature of this role in terms of pro- or antihypertrophic activity remains unclear and could be cell- and context-dependent. PGE2 may therefore not only be a passive consequence of prohypertrophic signaling during MSC chondrogenesis but may also either actively contribute to their endochondral misdifferentiation or attenuate chondrocyte hypertrophy. Defining this specific role of PGE2 for chondrocyte hypertrophy is important for understanding whether targeting COX/PGE2 can improve current MSC-based cartilage tissue engineering, how common COX-inhibiting pain medication can affect the outcome of MSC-based cartilage repair strategies, and could also have implications for the pain management of osteoarthritis patients.

In order to decipher the specific role of PGE2 for chondrocyte hypertrophy, here we aimed to uncover drivers of PGE2 secretion and its effects on hypertrophy during human MSC chondrogenesis. Stimulators of PGE2 release by chondrogenic MSC cultures were determined by inhibitor or withdrawal studies. The role of PGE2 for endochondral misdifferentiation was assessed by treatment with PGE2, the clinically applied non-steroidal anti-inflammatory drug celecoxib that inhibits COX-2, or the PGE2 receptor antagonists AH6809 or AH23848. Evaluation criteria were proteoglycan and type II collagen deposition as well as the expression of hypertrophy and osteogenesis markers and ALP enzyme activity.

## Results

### TGF-β, insulin, and endogenous BMP, WNT, and IHH activity jointly overcome PGE2 inhibition by dexamethasone

First, we documented the high PGE2 secretion of human bone marrow-derived MSCs in pellet culture with standard serum-free chondrogenic medium containing 10ng/mL TGF-β1, 6.25µg/mL insulin and 1µM dexamethasone. Histological analysis of day 42 samples revealed cells with a round chondrocyte-like morphology embedded in extracellular matrix rich in evenly distributed proteoglycans and type II collagen (Figure 1A), demonstrating successful chondrocyte formation. Consistent with our earlier findings [10], an ELISA assay detected 6.2±9.3ng/mL PGE2 (mean±SEM) in day 7 supernatants (Figure 1B). PGE2 levels increased significantly to 14.7±10.6ng/mL until day 21 (p=0.020) and then either stayed high (4 out of 11 independent donor populations) or declined again (7 out of 11).

**Figure 1.**
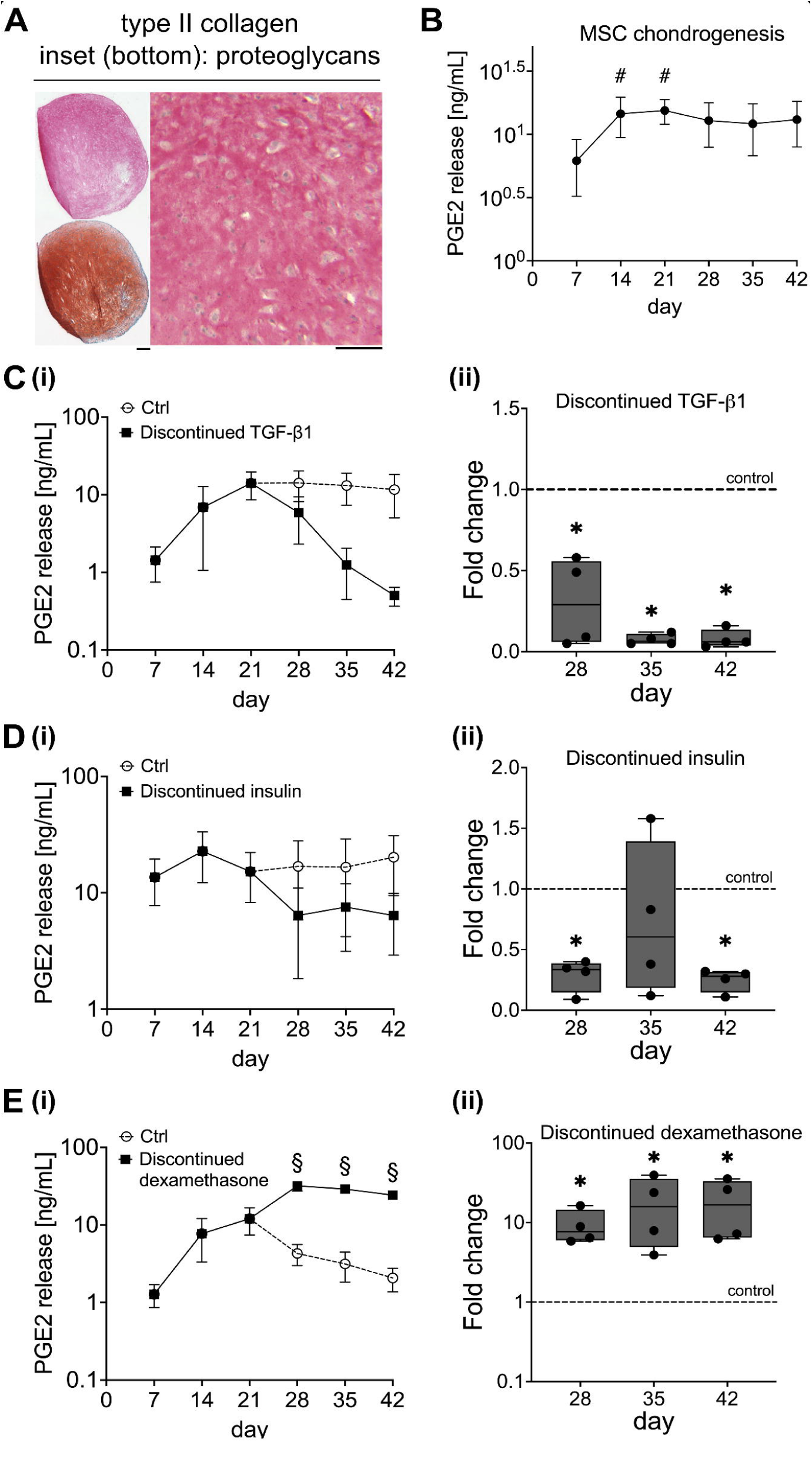
TGF-(31 and insulin drive while dexamethasone inhibits PGE2 release during MSC chondrogenesis. (A) Histological analysis of MSC differentiation. Paraffin microsections of day 42 samples were stained with Safranin O/Fast Green to visualize proteoglycan accumulation, and type II collagen was detected via immunohistochemistry (scale bar: 200µm, n=11). (B-E) Medium conditioned for 48 hours by differentiating MSC pellets was collected at weekly intervals and analyzed for PGE2 levels using ELISA (n=10-11). Controls received full chondrogenic medium, while in treatment groups, TGF-(31 (1Ong/ml), insulin (6.25µg/ml), or dexamethasone (0.1µM) were discontinued starting at day 21 as indicated (n=4). N designates the number of replicate experiments using independent MSC donor populations. Data points in the line diagrams represent mean±SEM. Box plots represent the interquartile range extending between the 25th and 75th percentiles, and lines inside the boxes represent the median. Whiskers extend to minimum and maximum values. The dashed lines indicate the control group set to 1. # p<0.05 vs. day 7, § p<0.05 vs. control, paired Student’s *t* test, *p<0.05 vs. control, Mann-Whitney U test.

To explore potential PGE2 drivers during MSC chondrogenesis, we first analysed the impact of the main chondro-instructive medium components by discontinuing the treatment with TGF-β1, insulin, or dexamethasone from day 21 on, which still allowed chondrogenic differentiation as documented by histological assessment (Supplemental Figure S1). Importantly, levels of secreted PGE2 decreased when TGF-β1 treatment was discontinued and reached levels as low as 0.5±0.3ng/mL on day 42 (Figure 1Ci). Setting the control to 1 to account for the known biological variability of MSCs revealed a significant PGE2 reduction by discontinued TGF-β1 at all assessed time points, reaching a mean reduction by 70% on day 28 (p=0.014) and 92% on days 35 and 42 (p=0.013, Figure 1Cii). PGE2 levels also declined when insulin treatment was discontinued, which reached a significant mean reduction by 90% at day 42 (p=0.014), yet remained a trend on day 35 (p=0.343, Figure 1D). By contrast, when the treatment with dexamethasone, a potent inhibitor of prostaglandin production, was discontinued, PGE2 release significantly increased almost 20-fold (day 42: p=0.014, Figure 1E). Altogether, these data identified TGF-β and insulin as drivers of PGE2 production during MSC in vitro chondrogenesis, acting against the inhibitor dexamethasone.

We next tested whether endogenous signaling also contributed to PGE2 release during MSC chondrogenesis and treated in adherence to previous studies [6–8, 33] with the BMP antagonist LDN-212854 (LDN-21, 500nM) from day 0 onward, with the WNT inhibitor IWP-2 (2μM) from day 14 on, or with the natural hedgehog antagonist PTHrP1-34 (daily 6-hour 2.5nM pulses) from day 7. Inhibitor functionality was confirmed by the ability of LDN-21 to reduce BMP-mediated SMAD1/5/9 phosphorylation, and of IWP-2 or pulsed PTHrP1-34 to inhibit ALP activity (Supplemental Figure S2). Consistent with previous data, all treatments allowed strong MSC chondrogenesis albeit LDN-21 slightly decreased while IWP-2 and pulsed PTHrP1-34 increased proteoglycan deposition according to histological assessment. Importantly, all three treatments – LDN-21, IWP-2, and pulsed PTHrP1-34 – strongly inhibited the upregulation of secreted PGE2 levels compared to controls (Figure 2). LDN-21 still allowed a slight upregulation of secreted PGE2 in all experiments and reached an 81% mean reduction at culture termination (p=0.037). In contrast, IWP-2 fully suppressed the upregulation of secreted PGE2 in all experiments and reached a mean reduction by 89% at day 42 (p=0.014), while pulsed PTHrP1-34 inhibited PGE2 upregulation in two out of three experiments and reached a mean peak reduction of 85% at day 28 (p=0.037), but only 61% at day 42 (p=0.037). To substantiate that the observed PGE2-inhibitory activity of PTHrP1-34 was attributed to hedgehog inhibition, we treated differentiating MSCs with the hedgehog receptor antagonist cyclopamine (10μM). Inhibiting as expected the expression of the hedgehog target gene *GLI1* (n=1, Supplemental Figure S3), cyclopamine allowed chondrogenesis according to histological assessment, and importantly, strongly reduced PGE2 levels. Overall, these data demonstrated that together with exogenous TGF-β and insulin, endogenous BMP, WNT, and hedgehog signaling stimulated PGE2 secretion during MSC chondrogenesis. Of note, two of these endogenous signals, i.e. WNT and hedgehog activity, are important prohypertrophic drivers of endochondral MSC misdifferentiation, and these data functionally connected chondrocyte hypertrophy with high PGE2 release.

**Figure 2.**
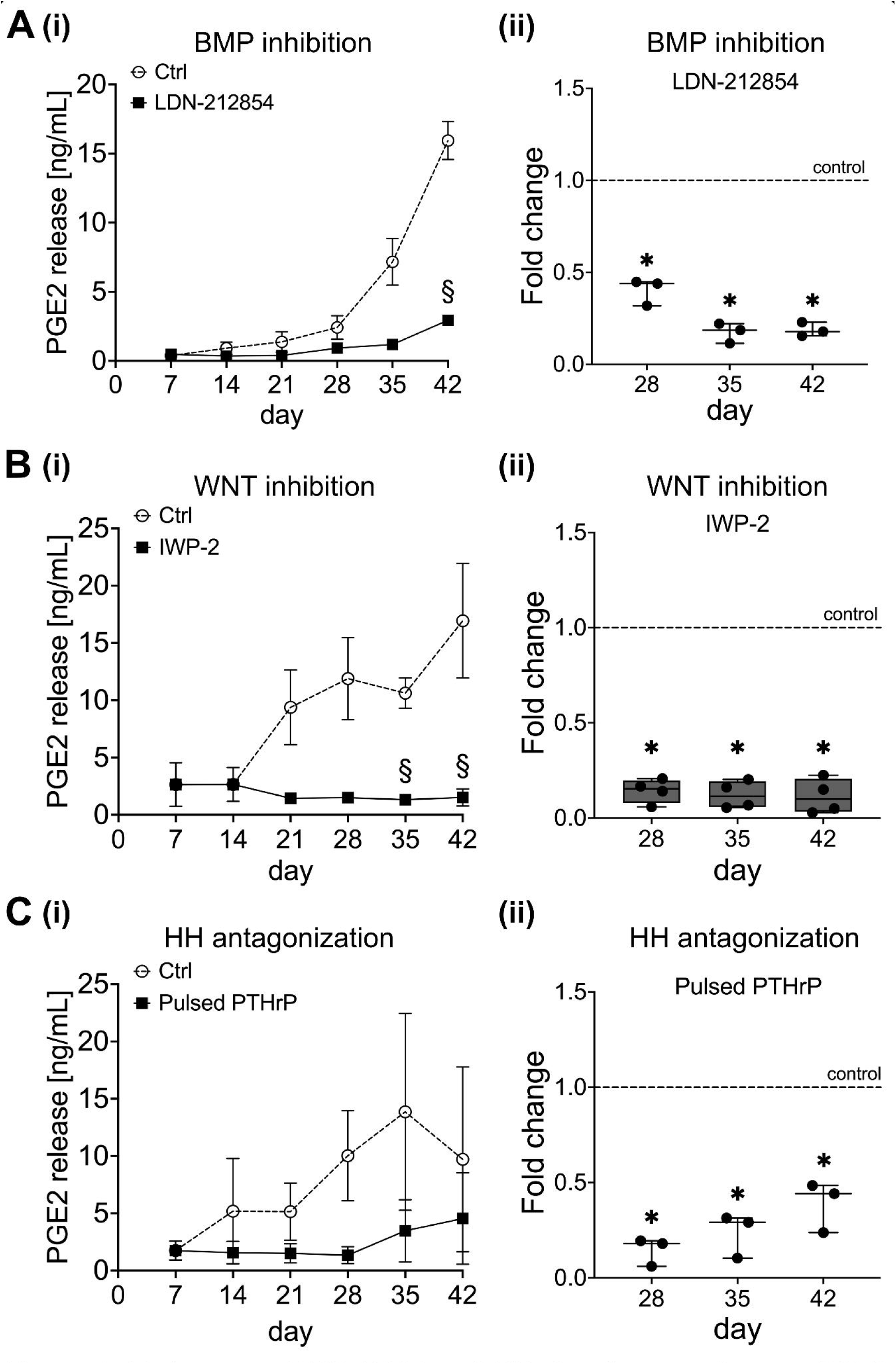
Endogenous BMP, WNT, and HH signaling contribute to PGE2 release during MSC chondrogenic differentiation. Cultures were treated as indicated with 500nM LDN-212854 from day O on (control: 0.02% DMSO, n=3), 2µM IWP-2 from day 14 on (control: 0.04% DMSO, n=4), or daily 6-hour pulses of 2.5nM PTHrP(1-34) from day 7 on (controls: daily medium exchange, n=3). Conditioned supernatants were assessed for PGE2 levels by ELISA. **N** designates the number of independent donor populations. Data points in the line diagrams represent mean±SEM, box plots are built as described in Figure 1. § p<0.05 vs. control, paired Student’s *t* test, * p<0.05 vs. control set to 1 indicated by a dashed line, Mann-Whitney U test.

### Unaltered chondrogenesis by prostaglandin E2 level modulation

To evaluate the possibility of autocrine PGE2 signaling in differentiating MSCs, we examined the expression of PGE2 receptors by interrogating our existing genome-wide expression data of MSCs at day 0 and day 28 of chondrogenesis [34] (Figure 3). While expression of both *PTGER1*/*EP1* and *PTGER3*/*EP3* was near or below the detection limit at both time points, *PTGER2*/EP2 and *PTGER4*/EP4 were highly expressed, which is consistent with a potential ability of differentiating MSCs to respond to PGE2. We chose the EP2-specific antagonist AH6809 and EP4 antagonists AH23848 in comparison with the COX-2-specific PGE2 synthesis inhibitor celecoxib for further experiments. Validation experiments demonstrated that 0.5µM celecoxib strongly suppressed secreted PGE2 levels during MSC chondrogenesis (Supplemental Figure S4A), while 10µM of AH6809 or AH23848 were capable to reduce the cAMP stimulation by 0.1µM PGE2 (Supplemental Figure S4B).

**Figure 3.**
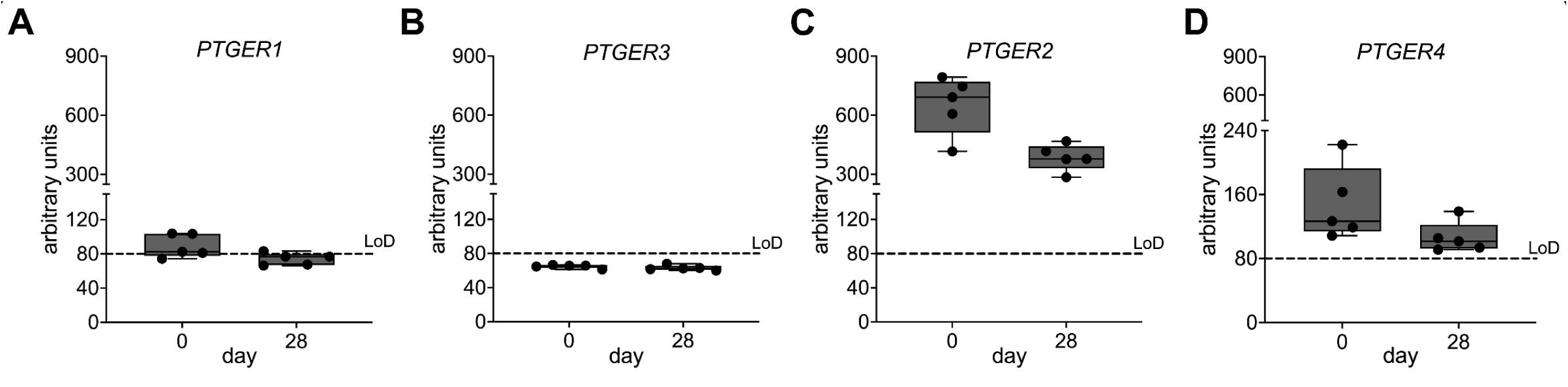
Prostaglandin E2 receptors 2 and 4 are expressed during MSC chondrogenesis. lllumina microarray data recorded in a previous study [34] was analyzed for gene expression levels of the four PGE2 receptors (n=5). The dashed lines represent the limit of detection (LoD), set at 80 arbitrary units, below which qPCR validation of target gene expression had failed in previous studies.

To next assess whether PGE2 plays an active role in MSC misdifferentiation and whether the EP2 and EP4 receptors mediate similar or different effects, we treated chondrogenic MSC cultures from day 0 on either with 1μM PGE2, or with 0.5μM celecoxib, 10μM AH6809 or AH23848, or 0.1% DMSO. After 6 weeks, strong and homogeneous deposition of cartilaginous matrix was observed histologically in all groups (Figure 4A). GAG/DNA quantification revealed an overall high proteoglycan amount in all groups with no significant differences caused by PGE2 level modulation (Supplemental Figure S5A). However, PGE2 appeared to decrease GAG/DNA levels in samples with lower proteoglycan content. Also, both celecoxib and AH23848 and to a lesser degree also AH6809 slightly decreased GAG/DNA levels, which became significant when DMSO controls were set to 1 (p=0.014, Figure 4B). Moreover, expression of the chondrocyte marker *COL2A1* rose significantly over time in all groups independently of PGE2 level modulation (Supplemental Figure S5B). PGE2 even slightly increased *COL2A1* gene expression by 21% compared to DMSO controls, which became significant when controls were set to 1 (p=0.005, Figure 4C). Again, this effect was not reversed by treatment with the inhibitors. The fibrocartilage marker type I collagen (*COL1A1*), commonly expressed during MSC chondrogenesis [35], was significantly decreased by PGE2 treatment by about 40% relative to the DMSO control (p=0.005, Figure 4D), while the inhibitors showed a donor-dependent trend to upregulate *COL1A1* mRNA (p=0.576, p=0.180, p=0.180). Overall, these data showed that PGE2 allows efficient differentiation of MSCs into chondrocytes and a strong deposition of the hallmark cartilage matrix components and slightly reduced the undesirable fibrocartilaginous phenotype.

**Figure 4.**
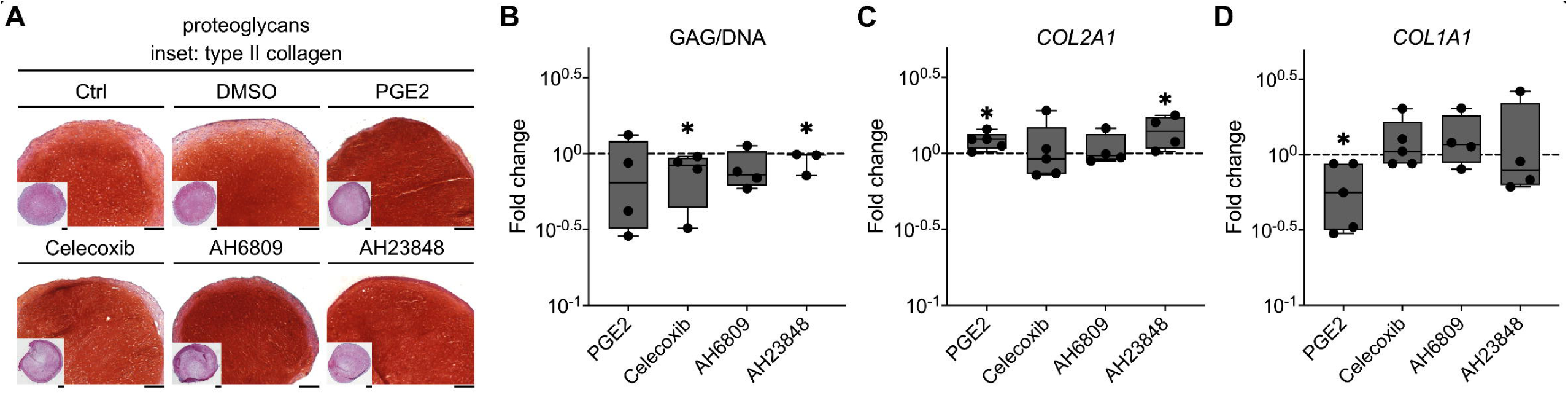
Effects of PGE2 level modification on chondrocyte differentiation and matrix deposition. MSC pellets were cultured in chondrogenic medium supplemented with PGE2 (1µM), the COX-2 inhibitor celecoxib (0.5µM), the EP2 and EP4 receptor antagonists AH6809 and AH23848 (10µM) respectively, or DMSO solvent (0.1%, n=4-5). Controls (Ctrl) received full standard chondrogenic medium. (A) Histological analysis of day 42 samples using Safranin O/Fast Green staining for proteoglycan visualization and immunohistochemistry for type II collagen detection (scale bar: 200µm). (B) Quantification of the proteoglycan content per pellet at day 42, assessed by DMMB assay, per total DNA amount assessed by the picogreen assay, related to values of DMSO controls (dashed line, n=3-4). (C, D) Gene expression of the chondrocyte marker *COL2A1* and the fibrocartilage marker *COL1A1* was quantified by qPCR at day 42 relative to DMSO controls (dashed line). *CPSF6, HPRT,* and *RPL13* were used as reference genes. Box plots are built as described in Figure 1, *p<0.05 vs. DMSO control, Mann-Whitney U test.

### Antihypertrophic activity of PGE2

Next, we assessed the effect of PGE2 on hypertrophy and osteogenesis markers. Interestingly, PGE2 treatment slightly but significantly reduced the expression of *COL10A1* by 18±3% (mean±SEM, p=0.005) compared to the DMSO control (Figure 5), along with a decreased expression of *MEF2C* in samples from 4 out of 5 MSC donors (reduction by 20±3%, p=0.095). Of note, the osteogenic markers *IBSP, SPP1*, and *ALPL,* appeared to be more strongly reduced by PGE2 by 63±7% (p=0.005), 77±3% (p=0.005), and 71±5% (p=0.005), respectively, compared to DMSO controls (Figure 5C-D, Supplemental Figure S5C). In support, PGE2 also significantly reduced the protein levels of the mineralizing enzyme ALP by 85±3% (p=0.005) at the end of differentiation according to an enzyme activity assay (Figure 5E-F). By contrast, inhibition of PGE2 synthesis using celecoxib significantly increased gene expression levels of *COL10A1* (178±23%, p=0.005), *MEF2C* (42±4%, p=0.005), *IBSP* (95±13%, p=0.005), *SPP1* (547±58%, p=0.005), and of *ALPL* (167±13%, p=0.095) by trend in 4 out of 5 MSC donors, compared to the controls. In line, ALP enzyme activity was significantly elevated by 64±10% (p=0.005) at culture termination (Figure 5E-F). Interestingly, neither the EP2 antagonist AH6809 nor the EP4 antagonist AH23848 fully reproduced the effects of celecoxib and opposed the PGE2 effect as reproducibly, suggesting a redundant function of these receptors. Altogether, this demonstrated that PGE2 is an autocrine antihypertrophic agonist during MSC chondrogenesis and seems to act via both EP2 and EP4 receptors with no apparent priority or effect-selectivity.

**Figure 5.**
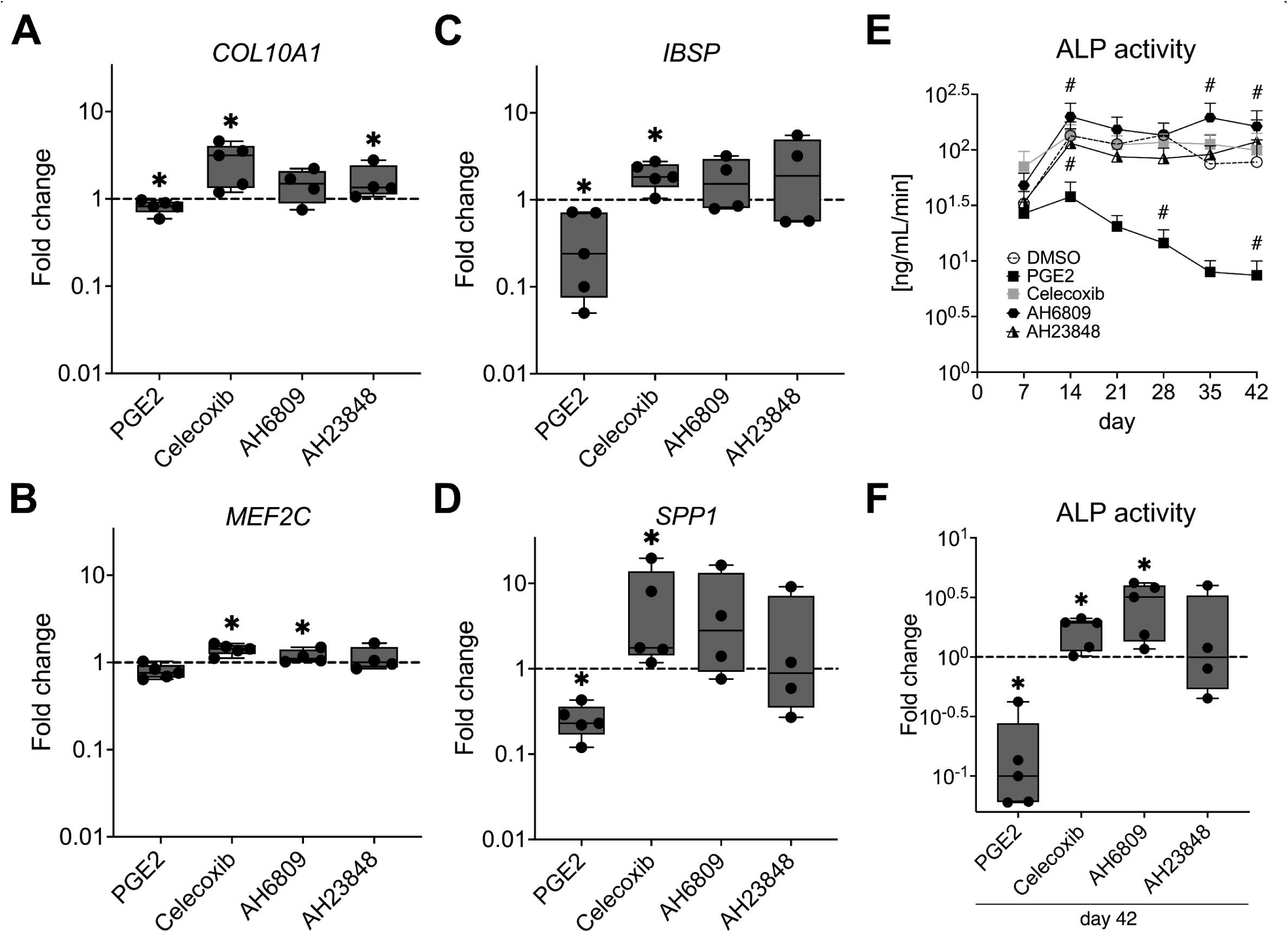
PGE2 exerts anti-hypertrophic and anti-osteogenic effects during MSC chondrogenesis. (A-D) MSC pellets were treated with chondrogenic medium containing PGE2 (1µM), celecoxib (O.5µM), AH68O9 and AH23848 (1OµM) respectively, or DMSO (0.1%) as control (n=4-5). Expression of the indicated genes was quantified using qPCR in day 42 samples. *CPSF6, HPRT,* and *RPL13* were used as reference genes. Data are referred to DMSO controls (dashed lines). (E-F) ALP enzyme activity in the supernatant. Data points in the line diagram represent mean±SEM, box plots are built as in Figure 1. # p<O.O5 vs. control, paired Student’s *t* test, *p<O.O5 vs. control set to 1, Mann-Whitney U test.

## Discussion

Prostaglandin E2 is a well-known inflammatory and immune modulatory mediator with a complex spectrum of activities in a multitude of diseases including osteoarthritis, in tissue regeneration, and in development. Intrigued by the conflicting evidence for pro- and antihypertrophic effects of PGE2 in various models of endochondral development, we here followed up on our previous discovery of highly elevated PGE2 release rates during MSC chondrogenesis when compared to articular chondrocytes (ACs) [10]. We here untangled for the first time that the main chondrogenic medium components all regulated PGE2 release from MSCs, whereby the COX-inhibiting dexamethasone was opposed by the stimulatory action of exogenous TGF-β and insulin. Since AC cultures under similar stimulation release very little PGE2, these exogenous signals may not be the main mediators of high PGE2 release from MSC cultures. Instead, the endogenous activity of BMP, WNT, and hedgehog signaling that are all well-known to be cell-autonomously overactivated in chondrogenic MSC compared with AC cultures [5, 6, 34], and were shown here to all stimulate PGE2 release from MSCs, appears to be responsible for the over 100-fold PGE2 overinduction [10]. Surprisingly, although the high PGE2 release was thus at least in part a consequence of the prohypertrophic signals that drive MSC misdifferentiation, PGE2 here turned out to be a broad inhibitor of chondrocyte hypertrophy, albeit its activity is obviously not sufficient to maintain MSCs stably in the chondral lineage. Thus, we here identified PGE2 as an endogenous autocrine antihypertrophic signal, whose inhibition would be detrimental for MSC-based engineered neocartilage by enhancing main aspects of the undesired process of endochondral bone formation.

As an immediate early gene, it is not surprising that *COX-2* and downstream PGE2 synthesis was affected by many signals. In line with our data, BMP, WNT, TGF-β, and insulin are all well-described PGE2 stimulators in various cell types [16–19, 36–38]. Only HH has rarely been attributed with this ability [39]. Interestingly, a previous study reported that the PGE2 response of chondrocytes to interleukin 1β is colocalized in the primary cilia together with mechanocoupling, hedgehog and WNT signaling, thus implying an interaction of these signals by physical colocation [40]. Of note, MSCs share this high PGE2 release with osteoarthritic chondrocytes [41] and it is tempting to assume that the here identified PGE2 stimulatory signals, i.e. TGF-β, BMP, hedgehog, and WNT, which have all been described to be overactivated also in osteoarthritis, may also contribute to PGE2 release from diseased chondrocytes.

A very important result in our study is that we revealed PGE2 to exert autocrine antihypertrophic effects on human hypertrophic chondrocytes. Since chondrocyte marker expression and cartilage matrix deposition remained largely unaffected by modulations of the PGE2 levels, this was not merely a reduction of the speed of chondrogenic differentiation. Instead, the here observed highly consistent opposite effects of increasing versus lowering PGE2 levels on all assessed hypertrophy and osteogenesis markers demonstrated that PGE2 is capable to partly decouple the undesired hypertrophic malformation from the desired chondrogenic differentiation. This antihypertrophic activity was well in line with the extensive studies by the O’Keefe group and others in chick and murine limb bud mesenchymal cells and growth plate chondrocytes [28–32], and with reports of a chondroprotective effect of EP2 and EP4 activity in osteoarthritic mouse models [30, 42]. Opposing results were reported primarily from the murine ATDC5 cell line, where PGE2 lowered and COX-2 inhibition enhanced *Col10a1* expression and ALP activity [20, 21]. Differences in the culture models, specifically the artificial monolayer induction of this cell line could be responsible for this discrepancy. In contrast to cell monocultures, the putative antihypertrophic PGE2 effect on chondrocytes would in complex in vivo conditions coincide with the well-known influence of PGE2 on vascularization, immune cell activity, and inflammation [12, 14, 24, 25, 43]. Therefore, the impeded bone development and regeneration in many COX-2/PGE2 loss-of-function studies is consistent with an inadequate PGE2-mediated stimulation of vascularization and osteoclast activation (amongst many others), which will critically interfere with bone remodelling despite potentially coinciding with enhanced chondrocyte maturation under relieved PGE2-mediated repression.

The antihypertrophic activity of PGE2 could potentially be harnessed to improve MSC-based cartilage tissue engineering and its instruction into the desired chondral instead of endochondral differentiation. Therefore, a reduction of the COX-inhibiting dexamethasone supply could be a promising alternative to PGE2 treatment but may also compromise chondrogenic induction. However, PGE2 did not fully suppress hypertrophy and osteogenic markers, even though we applied PGE2 in a supraphysiological concentration. We therefore did not follow up on the potential self-sustaining capability of the engineered neocartilage and its resistance to bone remodeling in vivo. Instead, it appears more promising to combine this new knowledge with previously discovered antihypertrophic treatments, i.e. with PTHrP pulses and WNT inhibition. However, careful dose titrations will be necessary because both treatments will inevitably reduce endogenous PGE2 production. Another important proof of concept would be to assess whether COX-2/PGE2 regulation are involved in instructing the chondral versus endochondral cell fate decision of MSCs in vivo that we are now able to instruct by adequately adjusting the sulfation rate of glycosaminoglycan hydrogels [44]. Since under chondral instruction, WNT and hedgehog signaling are not induced, and BMP signaling is lowered, we would expect very little PGE2 secretion; however, this remains to be investigated in a future study.

Beyond MSC-based cartilage tissue engineering, our results may also have important implications for osteoarthritis, for which MSC chondrogenesis is an interesting in vitro model. Accurately phenocopied osteoarthritis aspects of this model include the hypertrophic degeneration of chondrocytes, upregulation of matrix-degrading enzymes including MMP13 and ADAMTS4 [45], as well as proinflammatory signaling e.g. through NFκB [10], and here shown also high PGE2 release and the apparent EP2 and EP4 receptor portfolio [46, 47]. Importantly, COX-2 inhibiting medication of the non-steroidal anti-inflammatory drug (NSAID) class is widely used in osteoarthritis analgesia. Beneficial inhibition of inflammation, vascularization, and osteoclast activation may be hampered by the loss of the here demonstrated chondroprotective PGE2 effects. In line, EP4 activation was shown to reduce chondrocyte hypertrophy in a mouse osteoarthritis model [30], and importantly, first cohort and observational studies suggested that long-term use of COX inhibiting medication may accelerate osteoarthritis in patients [48, 49]. Thus, beneficial and detrimental effects of NSAIDs on the multiple osteoarthritis pathologies will need more careful consideration and further mechanistical investigations.

In conclusion, we here demonstrated for the first time that signals that can drive chondrocyte hypertrophy and include TGF-β and insulin together with cell-autonomous BMP, WNT, and hedgehog activity, stimulate PGE2 release of human MSC-derived chondrocytes. Beyond its paracrine effects, this PGE2 can autocrinely signal back to hypertrophic chondrocytes and dampen the prohypertrophic effects of the signals that drive its production. The broad antihypertrophic effects of PGE2 could in future be harnessed in the form of combinatorial treatments to improve the currently unstable cartilage tissue engineering approaches with MSCs. Moreover, the potential loss of this chondroprotective PGE2 activity as a side effect of NSAID administration in osteoarthritis patients should be followed up and COX-independent analgesics should be tested as alternative.

## Methods

### Isolation and culture of human bone marrow-derived MSCs

All procedures were approved by the local Ethics Committee for Human Experimentation of the Medical Faculty at Heidelberg University and followed the 1975 Helsinki Declaration in its latest version. Human MSCs were isolated with written informed consent from bone marrow of patients (age 33-83) undergoing total hip arthroplasty surgery. Mononuclear cells were enriched by a Ficoll-Paque™ PLUS (Cytiva, Freiburg, Germany) density gradient and cultured in 0.1% gelatin-coated culture flasks (37°C, 6% CO2) with expansion medium consisting of Dulbecco’s Modified Eagle’s Medium (DMEM), high glucose, 12.5% fetal bovine serum, 1% penicillin/streptomycin (Pen Strep, Gibco™, Thermo Fisher Scientific, Darmstadt, Germany), 2mM L-glutamine, 1% minimum essential medium non-essential amino acids, 1% 2-mercaptoethanol (all Gibco™, Thermo Fisher), and 4ng/mL fibroblast growth factor-2 (Active Bioscience, Hamburg, Germany). Non-adherent cells were washed off after 24 hours with phosphate-buffered saline (PBS) with the medium changed three times per week.

5x10^5^ passage 3 MSCs were pelleted and cultured for 42 days in chondrogenic medium consisting of DMEM high-glucose, 1% Pen Strep, 0.1μM dexamethasone, 0.17mM L-ascorbic acid 2-phosphate, 2mM sodium pyruvate, 0.35mM proline (all Sigma-Aldrich, Darmstadt, Germany), 1% ITS™+ Premix (Corning Life Sciences, New York City, USA), and 10ng/mL TGF-β1 (PeproTech, Darmstadt, Germany), supplemented as indicated with the BMP inhibitor LDN-212854 (500nM in dimethyl sulfoxide (DMSO), day 0-42; Sigma-Aldrich), the WNT signaling inhibitor IWP-2 (2μM in DMSO, day 21-42, Tocris Bioscience, Bristol, UK), the hedgehog inhibitor cyclopamine (10μM in DMSO, day 14-42; Tocris Bioscience), parathyroid hormone-related protein(1-34) (PTHrP1-34; 2.5nM in deionized water, day 7-42; Bachem, Weil am Rhein, Germany), PGE2 (1μM in DMSO, day 0-42; Cayman Chemical, Ann Arbor, MI, USA), the COX-2 inhibitor celecoxib (0.5μM in DMSO, day 0-42; LC Laboratories, Woburn, MA, USA), the EP2 receptor antagonist AH6809 (10μM in DMSO, day 0-42; Sigma-Aldrich) and the EP4 receptor antagonist AH23848 (10μM in DMSO, day 0-42; Apexbio Technology, Houston, TX, USA). When applicable, pellets were treated with DMSO as a solvent control. Medium was changed three times a week.

### PGE2 enzyme-linked immunosorbent assay (ELISA)

48-hour conditioned culture supernatants were collected at the designated time points and pooled from 4-5 pellets per treatment group and donor. A colorimetric competitive PGE2 ELISA (Enzo Life Sciences, Farmingdale, NY, USA) was performed according to the manufacturer’s instructions.

### Quantification of proteoglycan and DNA content

One pellet per group and donor was digested by 0.1mg/mL proteinase K (Thermo Fisher) in Tris hydrochloride (50mM, Carl Roth, Karlsruhe, Germany, 1mM CaCl2, Sigma-Aldrich) for 18 hours at 60°C under constant agitation. Samples were diluted in Tris/EDTA buffer (Invitrogen, Thermo Fisher Scientific) and mixed with 1,9-dimethyl-methylene blue solution (48μM, Thermo Fisher Scientific, in 40mM NaCl, 40mM glycine, both Carl Roth). The absorbance was measured at 530nm and referred to a chondroitin 6-sulfate standard curve.

The DNA content was quantified in proteinase K digested samples using the Quant-iT™-PicoGreen^®^ kit (Invitrogen) according to the manufacturer’s protocol. Obtained fluorescence values were referred to a λ-DNA-derived standard curve.

### Gene expression analysis

2-3 pellets per group and donor were pooled, polytronized and RNA was extracted using peqGOLD TriFast™ reagent (Peqlab, VWR, Avantor, Erlangen, Germany) according to the manufacturer’s protocol. 500ng total RNA were transcribed into complementary DNA using oligo-dT primers and OmniScript^®^ Reverse Transcription kit (QIAGEN, Hilden, Germany). Transcription levels of genes of interest were assessed by quantitative PCR (qPCR) using SYBR^®^ Green (Thermo Fisher) and a LightCycler^®^ R 96 instrument (Roche Diagnostics, Basel, Switzerland) using specific primer sequences (supplementary Table S1). Relative gene expression was calculated as 1.8^-ΔCt^, where ΔCt is the difference between the gene of interest Ct value and the arithmetic mean Ct value of the reference genes *CPSF6*, *HPRT*, and *RPL13*. qPCR products were quality controlled by agarose gel electrophoresis and melting curve analysis.

### ALP activity assay

Supernatants from 4-5 pellets per group were pooled and mixed with p-nitrophenyl phosphate substrate (10mg/mL in 0.1M glycine, pH 9.6, Carl Roth, 1mM MgCl2, and 1mM ZnCl2, both Sigma-Aldrich) and substrate conversion was measured after 90 min at 405nm versus 490nm. Sample values were referred to a p-nitrophenol (Sigma-Aldrich) standard curve and calculated as ng/mL/min.

### Histology and immunohistochemistry

On day 42 of chondrogenic differentiation, MSC pellets were fixed in 4% paraformaldehyde (in PBS, pH 7.4), dehydrated in graded propan-2-ol, and embedded in paraffin. 5μm microsections were deparaffinized, stained with Safranin O (0.2% w/v in 1% acetic acid, Sigma-Aldrich) for proteoglycan detection, and counterstained with Fast Green (0.04% w/v in 1% acetic acid, Merck, Darmstadt, Germany).

Type II collagen immunostaining was conducted as previously established. In brief, deparaffinized microsections were rehydrated and incubated with hyaluronidase (4mg/mL in PBS, pH 5.5) followed by pronase (1mg/mL in PBS, pH 7.4, both Roche Diagnostics, Mannheim, Germany). After blocking with 5% bovine serum albumin sections were incubated with a mouse anti-human type II collagen antibody (1:1000, clone II-4C11, MP Biomedicals, Santa Ana, CA, USA), followed by an ALP-coupled goat anti-mouse immunoglobulin G secondary antibody (ImmunoLogic, WellMed, MS Arnhem, Netherlands) and ImmPACT^®^ Vector^®^ Red Substrate (Vector Laboratories, Newark, CA, USA) visualization.

### Statistics

The number of independent biological replicates of each experiment is indicated in the figure captions. Data was analyzed using SPSS Statistics (Version 29.0.0.0, IBM, Armonk, NY, USA). Mann-Whitney U test was used to assess differences presented in box plots and a paired Student’s *t* test tor time course experiments with a p-value <0.05 considered as statistically significant.

## CRediT authorship contribution statement

**S. Schmidt:** Investigation, Methodology, Formal analysis, Data curation, Visualization, Writing – original draft, Writing – review & editing. **F.A.M. Klampfleuthner:** Investigation, Methodology, Writing – review & editing. **T. Renkawitz:** Formal, analysis, Resources, Funding acquisition. **S. Diederichs:** Conceptualization, Formal analysis, Project administration, Supervision, Visualization, Writing – original draft, Writing – review & editing, Resources, Funding acquisition. All authors have read and agreed to the submitted version of the manuscript. **S. Schmidt** and **S. Diederichs** take full responsibility for the integrity of the work as a whole.

## Acknowledgements

The authors thank Prof. Dr. Wiltrud Richter for support in study conceptualization, Dr. Jennifer Fischer for experimental support, the clinicians of the Orthopaedic University Hospital Heidelberg for supplying primary patient material, and the patients for donating bone marrow specimens.

## Research ethics and patient consent

Each procedure was approved by the local Ethics Committee for Human Experimentation of the Medical Faculty of the University of Heidelberg and followed the Helsinki

Declaration of 1975, as revised in 2000. Human MSCs were isolated from the bone marrow after written informed consent.

## Data availability statement

The authors emphasize that none of the data generated as part of this study will be kept restrictive and are available upon reasonable request.

