## Supplementary figures and images for "Cause and chondroprotective effects of prostaglandin E2 secretion during mesenchymal stromal cell chondrogenesis"

### Supplemental Figure S1

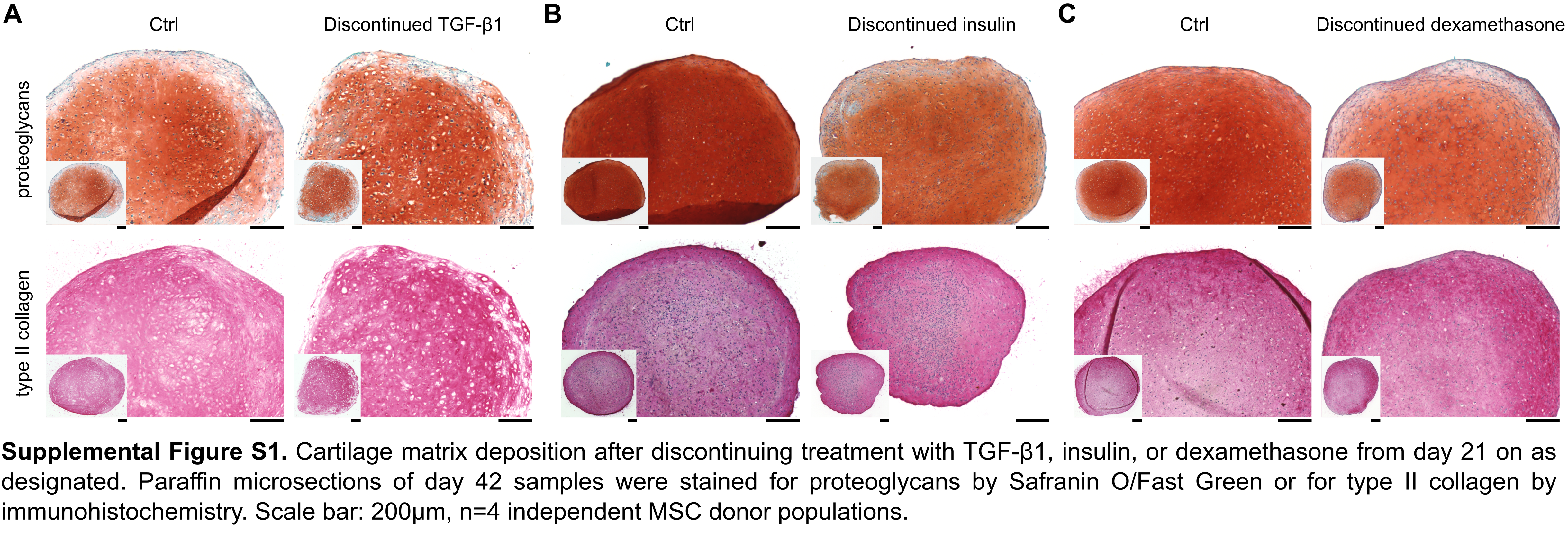

### Supplemental Figure S3

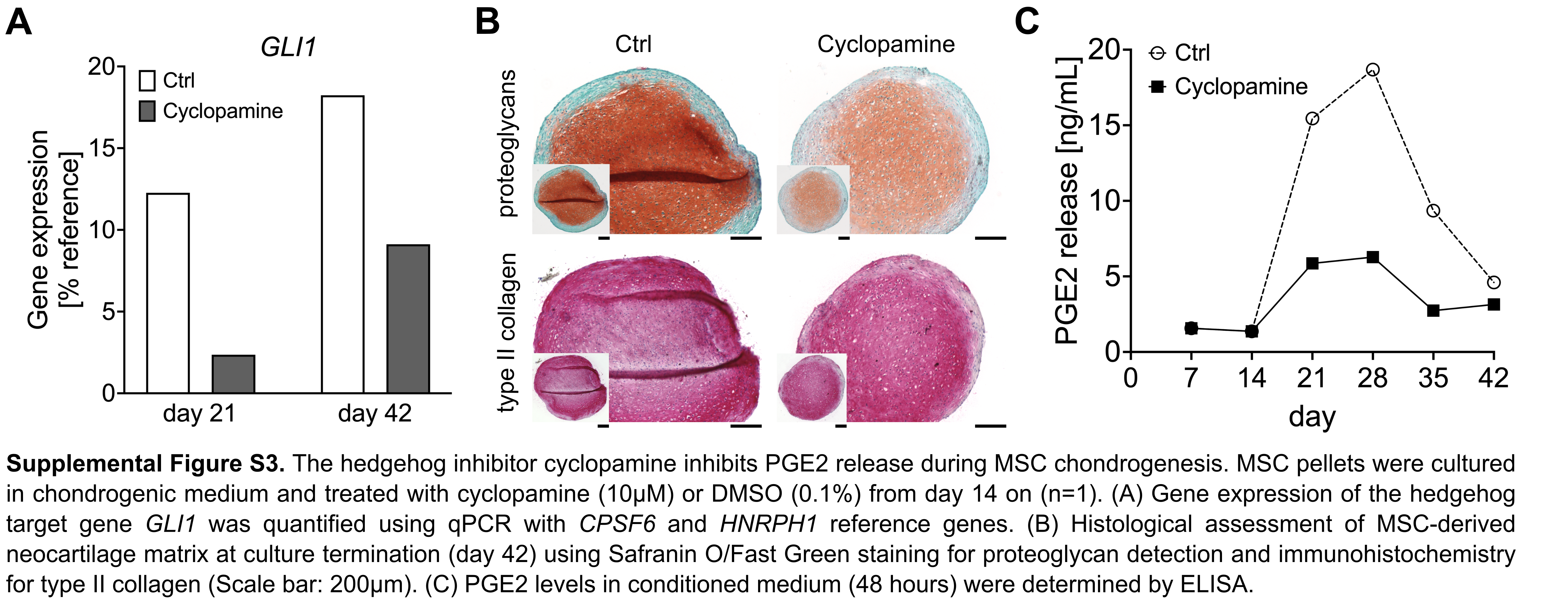

### Supplemental Figure S4

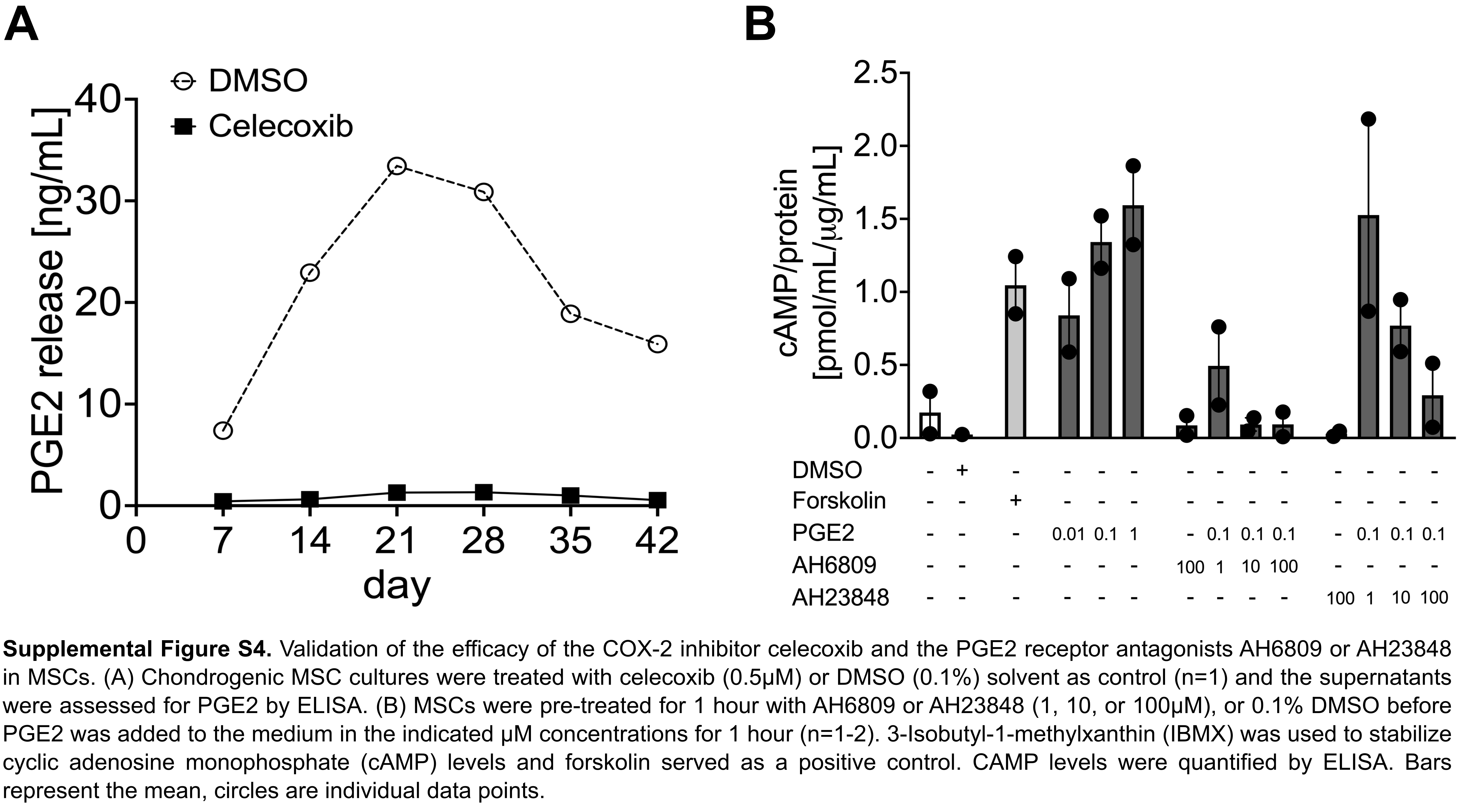

### Supplemental Figure S5

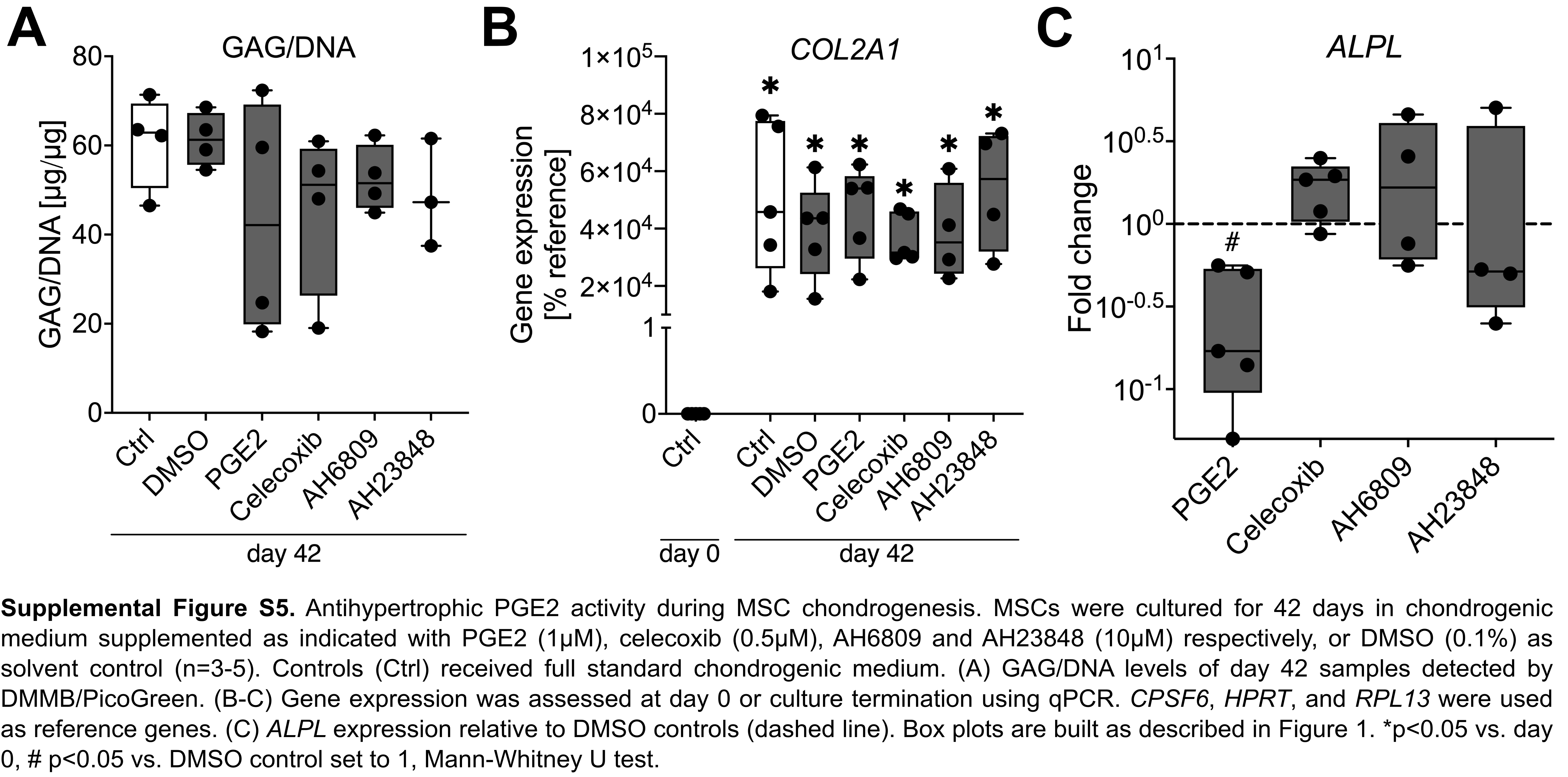
