## Supplemental Table S1 for "Cause and chondroprotective effects of prostaglandin E2 secretion during mesenchymal stromal cell chondrogenesis"

**Supplemental Table S1.** List of primer sequences used for qPCR in alphabetical order.

| Gen | Forward | Reverse |
|---|---|---|
| *ALPL* | 5´-CACCAACGTGGCTAAGAATG-3´ | 5´TCAGCTGGATGGCCACATC- 3´ |
| *COL1A1* | 5´-GATGCCAATGTGGTTCGTGA3´ | 5´-ATCTCCAGCCTGGTCTCCTC-3´ |
| *COL10A1* | 5´-TTTACGCTGAACGATACCAAA-3´ | 5´-TTGCTCTCCTCTTACTGCTAT-3´ |
| *COL2A1* | 5´-TGGCCTGAGACAGCATGAC-3´ | 5´-AGTGTTGGGAGCCAGATTGT-3´ |
| *CPSF6* | 5´-AAGATTGCCTTCATGGAATTGAG-3´ | 5´-TCGTGATCTACTATGGTCCCTCTCT-3´ |
| *HPRT* | 5´-AAGGGTGTTTATTCCTCATGGA-3´ | 5´-CCTCCCATCTCCTTCATCAC-3´ |
| *IBSP* | 5´-CAGGGCAGTAGTGACTCATCC-3´ | 5´-TCGATTCTTCATTGTTTTCTCCT-3´ |
| *MEF2C* | 5´-GTATGGCAATCCCCGAAACT-3´ | 5´-ATCGTATTCTTGCTGCCTGG -3´ |
| *RPL13* | 5´-CATTTCTGGCAATTTCTACAG-3´ | 5´-CAGGCAACGCATGAGGAAT-3´ |
| *SPP1* | 5´-GCTTGGTTGTCAGCAGCA-3´ | 5´-TGCAATTCTCATGGTAGTGAGTTT-3´ |
